# Old worms, new tricks: dynamical instability explains late-life rejuvenation in *C. elegans*

**DOI:** 10.64898/2026.05.01.722260

**Authors:** Djakim Latumalea, Adrian Molière, Peter Fedichev, Collin Y. Ewald, Jan Gruber

**Affiliations:** Department of Biochemistry, Yong Loo Lin School of Medicine, National University of Singapore, Singapore; Department of Health Sciences and Technology, ETH Zürich, Switzerland; Gero PTE. LTD., 133 Cecil Street 14-01 Keck Seng Tower, Singapore 069535

**Keywords:** physics of aging, Langevin model, rejuvenation, longevity, *daf-2*, temperature, proteohomeostasis

## Abstract

How is it possible to double the lifespan of an organism already close to death? Many biological theories of aging fail to explain this phenomenon. At the Physics of Aging workshop, we presented and discussed late-life lifespan extension in *Caenorhabditis elegans* to illustrate how a simple stochastic dynamical systems model can account for dramatic geriatric interventions. We build on a Langevin-type instability framework in which aging is a manifestation of dynamical instability–a scenario where stochastic fluctuations amplify over time, driving the system toward a failure threshold at which death occurs as a first-passage event. The instability rate *α* (equivalently, the inverse of the mortality-rate doubling time) quantifies the speed of this divergence: a larger *α* means faster exponential growth of *z*, a steeper Gompertz slope, and a shorter lifespan. The failure threshold *z*_max_ *≈α/g*, where *g* is the strength of nonlinear feedback, marks the point beyond which the system diverges irreversibly—physiologically, the saturation of metabolic and regulatory capacity. Within this dynamical-systems framework, auxin-induced degradation of the insulin/IGF-1 receptor DAF-2 in very old animals is naturally interpreted as a late shift in stability parameters that nearly doubles remaining lifespan without resetting accumulated structural damage. This interpretation reconciles the persistence of many senescent pathologies with restored proteostasis and stress resilience, and it shows that targeting the dynamical instability of the regulatory network–rather than reversing damage—can strongly reshape survival trajectories in unstable animals. More broadly, our work exemplifies how physics-inspired low-dimensional stochastic models can capture key features of aging, and we hope it will inspire more collaborations between biologists and physicists to work on late-life interventions.

## INTRODUCTION

An ancient dream is to age gracefully and then magically rejuvenate the body to continue living without the pathologies and diseases of aging. It is widely assumed that such rejuvenation of an old body is impossible; although this appears to hold true for humans so far, it is not necessarily the case across all species. Indeed, we have shown that the nematode *C. elegans* is capable of something reminiscent of this phenomenon. The multicellular model organism *C. elegans* typically has a lifespan of around 15 to 25 days [1]. Remarkably, treatment in geriatric nematodes older than 20 days can still double their lifespan, yielding an additional 20 days of survival [2]—a surprising finding that has been reproduced by other laboratories [3]. This was achieved using a novel genetically engineered protein degradation system, auxin-induced degradation, which enables precise, temporally controlled degradation of the insulin/IGF-1 receptor DAF-2 at any age [2].

Mutations in the insulin/IGF-1 receptor DAF-2, as well as knockdown of *daf-2* by RNA interference (RNAi), can double the lifespan of *C. elegans* to more than 60 days [4, 5]. However, *daf-2* mutations alter insulin/IGF-1 receptor signaling from birth, and the latest age at which *daf-2* knockdown by RNAi is able to extend lifespan is day 6 of adulthood, when *C. elegans* is still repro-ducing [6]. Historically, this led to the conclusion that reducing insulin/IGF-1 receptor signaling during development and early adulthood can increase lifespan, but not beyond the reproductive period [6].

This interpretation aligns with a damage-accumulation view of aging [7]. If aging were primarily driven by the progressive accumulation of irreversible molecular damage, then interventions that merely slow the formation of damage should show diminishing returns when initiated late in life. By the time treatment begins, a substantial fraction of critical lesions—including misfolded proteins, crosslinked extracellular matrix components, mitochondrial dysfunction, and genomic instability–would already be present. In such a framework, true rejuvenation would be at best partial, if not impossible, because it would require reversal of all forms of entropic damage. Even partial removal or repair of accumulated macromolecular damage would be energetically costly, mechanistically complex, and unlikely to occur rapidly or completely in a post-reproductive organism. One would therefore predict that late-life interventions could, at best, modestly slow further decline, but not produce large, immediate extensions of remaining lifespan. The reported decline in the efficacy of late-life *daf-2* RNAi has therefore historically been interpreted as evidence that aging in *C. elegans* progresses beyond a point at which modulating damage accumulation can meaningfully alter survival [6].

However, the expression of various RNAs that regulate the RNAi machinery also changes with age [8], suggesting that age-dependent variation in *daf-2* RNAi knock-down efficiency, rather than diminished biological return from *daf-2* knock-down itself, may be responsible. This distinction raises a critical question: does the reduced benefit truly indicate that accumulated damage has rendered late-life intervention ineffective, or does it instead reflect a decline in the efficacy of the RNAi method itself?

## THE DAF-2 DEGRON SYSTEM

To address this question, we used a plant-derived genetic tool adapted for *C. elegans*: the auxin-induced degradation system [9]. We tagged the DAF-2 receptor with a small degron tag and introduced the plant F-box protein TIR1, a component of an SCF E3 ubiquitin ligase, into all *C. elegans* cells [2]. Under standard conditions, this degron-tagged DAF-2 is not degraded, functions normally, and animals appear wild-type. Upon auxin treatment, however, the degron-tagged DAF-2 is degraded within 30 minutes. Crucially, degradation is reversible: auxin withdrawal restores DAF-2 levels, enabling precise temporal control of the receptor at any age [2].

We used this system to test whether DAF-2 degradation initiated in geriatric animals, well beyond the window accessible to RNAi, could still extend lifespan. We showed that even well past the reproductive age, up to day 21 of adulthood, when 75% of the population had already died and when control-treated *C. elegans* only lived 4 additional days on average, auxin-mediated degradation of DAF-2 extended survival by an average of 26 days. In other words, total lifespan could still be doubled even when intervention was initiated near the end of life [2]. This remarkable finding has been reproduced by other laboratories, including the Gems Lab [3].

This finding raised the core question: *how is it possible to double the lifespan of a multicellular organism so close to the end of its normal lifespan?* In collaboration with several laboratories, we began to dissect which age-related pathologies are slowed, halted, or even rejuvenated when lifespan is extended at geriatric ages in *C. elegans* [3]. We found that, despite the robust lifespan extension, some age-related and morphological senescent pathologies (pharyngeal degeneration, gonadal atrophy, yolk pooling, and uterine tumors) are not restored or reversed [3], suggesting that these forms of “damage” are not primary determinants of late-life longevity in *C. elegans*. By contrast, age-related deterioration of the exoskeleton (cuticle)—which is composed of collagen and extracellular matrix proteins—is halted [3]. The decline in healthspan, including motility, is slowed [3]. Interestingly, resilience to heat and osmotic stress is also significantly restored [3]. Furthermore, age-related endoge-nous protein aggregation is resolved following late-life longevity intervention [3].

Taken together, these observations are difficult to reconcile with the damage-accumulation view of aging. If late-life mortality were primarily determined by the burden of irreversible molecular and structural lesions, then interventions initiated after substantial deterioration should yield only modest gains, and certainly should not restore remaining lifespan to the same upper limit achievable when treatment begins in young animals. Notably, the late-life *daf-2* degron intervention restores lifespan to the same maximal limit observed when the pathway is suppressed from early adulthood [2]. This occurs despite the persistence of extensive age-associated damage and structural pathology. The persistence of many age-associated pathologies despite dramatic lifespan extension further suggests that such lesions are not the principal determinants of survival risk and lifespan. This points to a fundamentally different determinant of late-life survival risk, one governed by the dynamical state of physiological regulation rather than the cumulative burden of structural damage.

## LANGEVIN INSTABILITY MODEL OF AGING

We therefore ask what dynamical property of the organism could account for both the late-life efficacy of DAF-2 degradation and the selective reversibility of age-related pathologies. A natural candidate is intrinsic instability—the possibility that *C. elegans*, and other short-lived organisms, operate in a regime where physiological trajectories diverge without restoring forces [10] (Fig. 1), extended in Fedichev & Gruber [11].

**FIG. 1.**
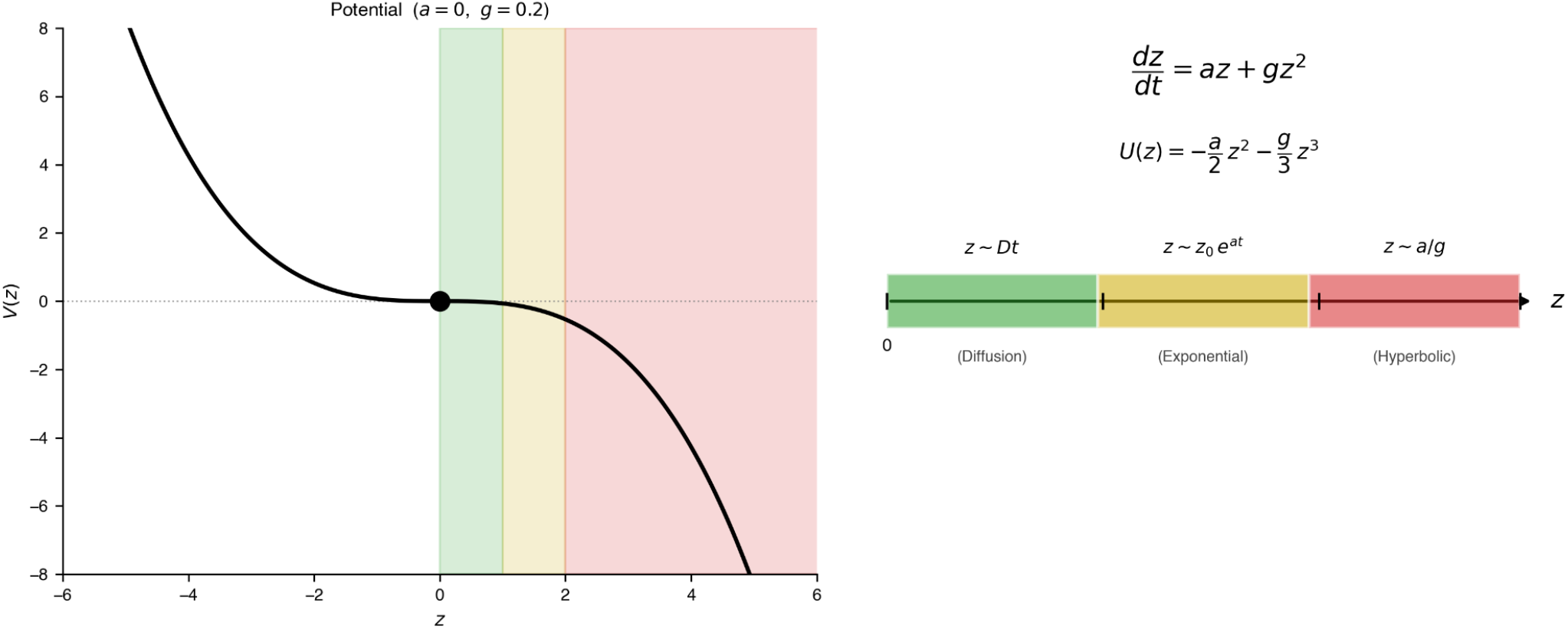
Instability dynamics of *C. elegans*. *Left* : Potential for the unstable case *α >* 0, the intrinsically unstable case in which linear stability (parameter *α*) drives exponential divergence from birth. Coloured bands delineate the three dynamical regimes encountered as a trajectory moves from *z* = 0 to *Z*: diffusion (green, *z ≈Dt*), where noise *D* dominates near the origin; exponential (yellow, *z ≈ z*_0_*e*^*αt*^), where the linear drift term drives growth; and hyperbolic (red, *z ≈ α/g*), where the nonlinear *gz*^2^ term accelerates divergence toward the death threshold. *Right* : Schematic summary of the same three regimes along the *z*-axis, together with the governing Langevin equation and potential.

Two features of this model are key to understanding the effects of late-life interventions. As we proposed in Fedichev & Gruber [11], aging manifests itself as a linear accumulation of irreversible (entropic) damage that affects the functional and stability properties of key physiological systems. In short-lived organisms such as *C. elegans*, the gradual entropic changes have little effect since the dominant aging signature is driven by dynamics of an inherently unstable mode—a cluster of tightly correlated features characterized by a single mode variable *z*—driving the exponential deviations of the physiological state from the youthful norm and, ultimately, to the death of the organism (Fig. 1).

Since aging is slow, the kinetics of aging follow the first-order stochastic Langevin equation. The key property is that the rate of change of *z* depends only on the current value of *z* (the Markov property). Although entropic damage *Z* accumulates throughout life, gradually eroding the stability parameter *ε*_0_, its effect is slow compared to the fast exponential divergence of *z* in the unstable regime. Mortality dynamics are therefore governed by the current value of *z* and the instantaneous parameters, not by the accumulated history of damage (Fig. 2, compare the two zoom-in boxes).

The remaining lifespan depends on the proximity to the failure threshold, *z*_max_ = *α/g*, where *α* is the instability (divergence) rate and *g* is the nonlinear feed-back strength. Below *z*_max_, the exponential regime dominates; beyond *z*_max_, nonlinear acceleration produces the hyperbolic (runaway) regime. Death is modeled as a first-passage event: the stochastic time at which *z* first reaches *z*_max_. Because noise causes different individuals to reach this threshold at different times, a distribution of lifes-pans is produced even from identical initial conditions (see [10] for more details). Representative trajectories illustrating this behavior are shown in Fig. 2, and the full dynamical evolution of the system is provided in Supplementary Video 1.

**FIG. 2.**
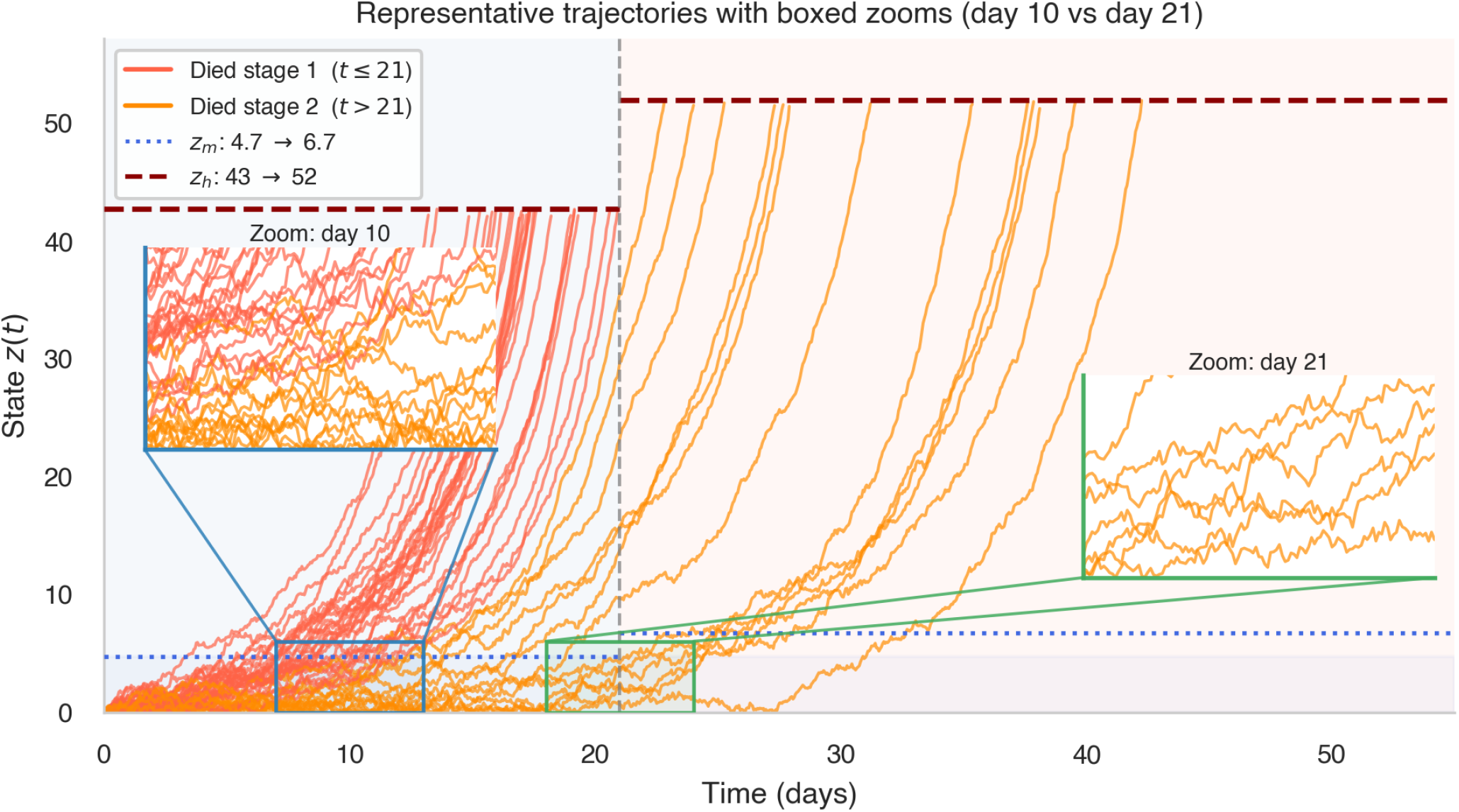
Trajectories reveal the absence of explicit time dependence. Representative stochastic trajectories *z*(*t*) from the Langevin model are shown for *C. elegans* that die before intervention (tomato), or die after intervention (dark orange). The vertical dashed line marks the intervention time; dotted horizontal lines indicate the drift-to-exponential level *z*_*m*_, and dashed horizontal lines denote the exponential-to-hyperbolic transition level *z*_*h*_. Boxed regions and corresponding insets compare dynamics at day 10 (early phase) and day 21 (after intervention). Despite the difference in chronological time, the local fluctuation structure and dispersion of trajectories within the low-state regime are indistinguishable. This reflects the Markov (i.e., memoryless) nature of the dynamics: future evolution depends only on the current state *z*(*t*), not on elapsed time. Late-life individuals occupying the same region of state space are dynamically equivalent to earlier ones, indicating that mortality risk is determined by instantaneous state rather than explicit time dependence.

One consequence of this model is that interventions modifying key dynamic parameters can remain effective even late in life. Let us follow how the effects of late-life DAF intervention naturally emerge from the model. Simulated Kaplan–Meier survival curves generated from the stochastic two-stage Langevin model reproduce the characteristic lifespan extension observed experimentally (Fig. 3A). In our simulations, we do not alter the noise power *D*_0_; the magnitude of stochastic fluctuations remains unchanged throughout life, both before and after the intervention. Instead, we modify only the dynamics by reducing the instability parameters. The fluctuations act with the same strength, but the system becomes less unstable: trajectories that would previously escalate rapidly into runaway hyperbolic failure remain in the deterministic dynamical (exponential) regime for longer (Fig. 2). Because the system is Markovian (i.e., memoryless), changes to the dynamics immediately increase the expected time to reach the hyperbolic failure boundary, even when intervention occurs late in life, and no damage removal or rejuvenation is required. The physiological state is not reset, and structural pathologies persist, yet lifespan is extended. Lifespan extension, therefore, does not require morphological rejuvenation, but only a shift in dynamical instability parameters. This interpretation aligns with experimental observations (Fig. 3B). Irreversible structural pathologies persist, consistent with the irreversibility of prior trajectories. Yet stress resistance improves, and protein aggregates resolve, suggesting that proteostasis stabilizes network dynamics. Restoring protein homeostasis does not rewind aging; it shifts the system farther from the tipping point.

**FIG. 3.**
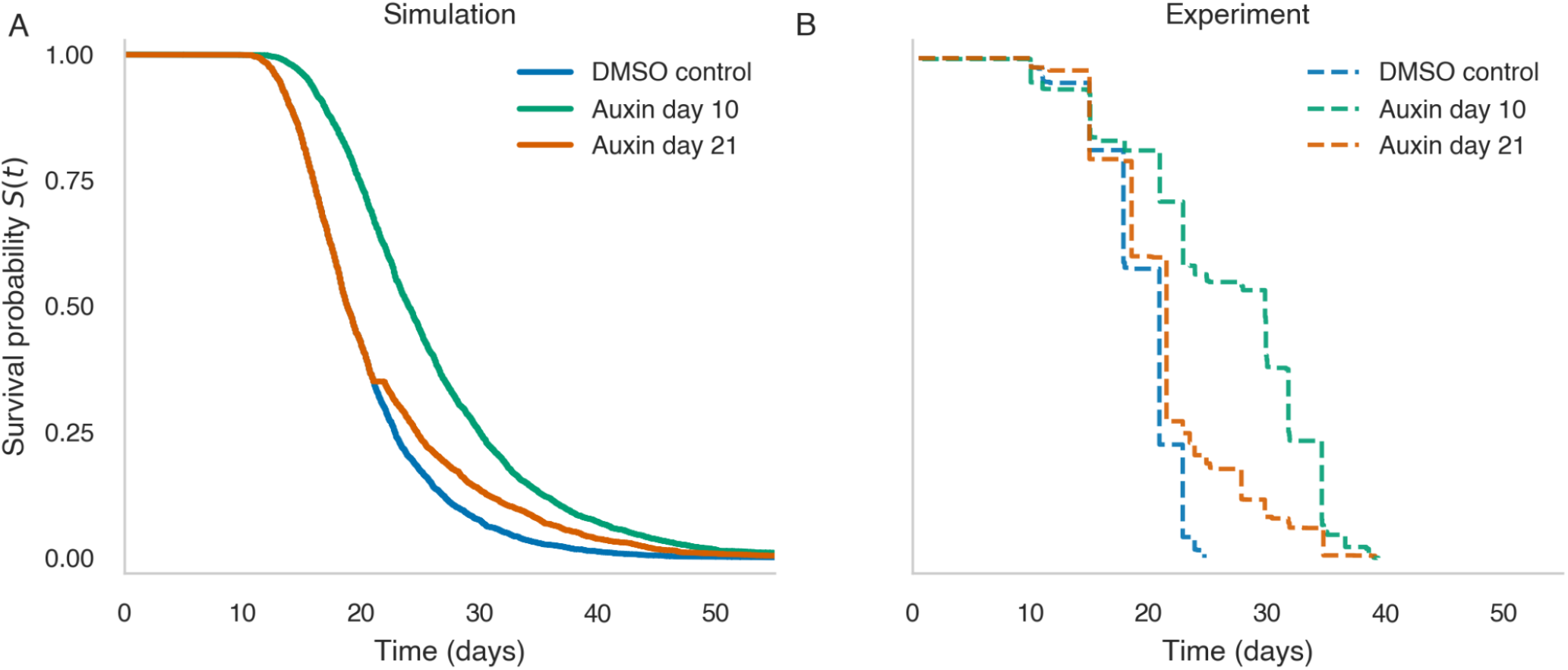
Simulated and experimental survival dynamics under early and late intervention. (A) Kaplan–Meier survival curves generated from the stochastic two-stage Langevin model. DMSO controls (blue) are compared with cohorts receiving intervention at day 10 (green) or day 21 (orange). (B) Experimental survival curves for the corresponding conditions (dashed lines), plotted on the same time scale.

## EVOLUTIONARY CONTEXT

To understand these observed phenomena, it is important to consider the life-history strategy of *C. elegans* in an evolutionary context. Two major variables governing the evolution of longevity are the age at sexual maturity and age-dependent fecundity schedules [12]. For reproduction to occur, an individual’s genome must be passed to the next generation; until then, selection pres-sure is maximal to keep the organism in optimal condition. With the onset of reproduction, the contribution of later age groups to the future gene pool diminishes, and thus the force of natural selection to maintain physiological integrity declines (Selection Shadow) [12]. Humans can and do reproduce for decades after reaching maturity, and therefore must have evolved mechanisms to ensure stability of their final body plan. This differs from animals that reproduce only once or within a very short window: systemic stability is no longer required, and the soma quickly becomes disposable. Aging driven by mechanisms that can be seen as a continuation of development, such as hyperfunction, would be expected to dominate in such unstable organisms [13, 14]. This is especially true when the post-reproductive adult provides no benefit to their offspring’s survival, or may even be detrimental. In *C. elegans*, it has been hypothesized that the death of the post-reproductive adult may even be advantageous, with nematodes actively converting their biomass into yolk for their offspring before dying and thereby making all their nutrients available to the next generation [15, 16]. In such a scenario, there may even be active pressure against the evolution of a stable post-reproductive system, which might explain why lifespan-extending interventions are so abundant in *C. elegans* [13]. Taken together, the evolved life history strategy of *C. elegans* is consistent with the idea that systemic instability is a core feature of *C. elegans* aging.

Importantly, this instability holds for *C. elegans* during normal development when resources are abundant. However, the dauer stage is an alternative developmental endpoint that has evolved to be stable over time, allowing survival through periods of scarcity until conditions are sufficient for reproduction. In that sense, *C. elegans* has both an unstable and a stable physiological mode. This explains that interventions acting through dauer signaling, like DAF-2 knockdown, are particularly potent at extending lifespan: *C. elegans* has an inbuilt stability program that can enhance stability even when activated outside its usual activation window, for example, at old age [17].

This physics-based Langevin model of aging is well aligned with other theories and frameworks of aging. The notion that short-lived animals with a short reproductive period, such as *C. elegans*, are inherently unstable is consistent with developmental theories that postulate aging as a quasi-programmatic continuation of development [18, 19]. A lack of stability after maturity, marked by continuing quasi-programs, is conceptually equivalent to instability characterized by high autocorrelation; the future state of the organism is expected to be derivable from the previous one. On the other hand, theories of deleterious accumulation of damage [20] are expected to apply more strongly to species that have evolved a stable period following maturation, marked by low autocorrelation. This suggests a continuum between stable and unstable species with potentially fundamentally different underlying mechanisms of aging. Another interesting interaction with the Langevin stability model concerns the framework proposed by Pio-Lopez et al. [21], which posits that aging is primarily a result of a loss of cellular goal-directedness following the completion of the developmental program. Without an overarching goal, systems are inherently unstable and prone to deterioration. Through this lens, the restoration of stability following DAF-2 auxin-induced degradation could be seen as the reestablishment of a physiological goal, driving the system in a programmed manner towards a state closer to the stable dauer stage.

## REFLECTIONS ON THE CORE QUESTIONS RAISED AT THE “PHYSICS OF AGING” WORKSHOP

The central question of our presented work was *“How is it possible to double the lifespan of a multicellular organism at the end of its lifespan?”* Classical theories of aging, such as damage accumulation, antagonistic pleiotropy, and hyperfunction, fall short in explaining the magnitude of late-life increase in lifespan. Thus a more suitable model was required to explain our experimental data. We examined a novel physical model of aging—a Langevin-inspired model based on non-equilibrium thermodynamics and statistical mechanics. We propose that in the model organism *C. elegans*, late-life mortality appears to be governed less by the cumulative burden of entropic damage and more by the dynamics of an intrinsically unstable physiological mode characterized by resilience, stochastic fluctuations, and proximity to a failure threshold. Thus the survival risk is determined by the organisms instantaneous dynamical state and stability parameters rather than by an irreversible record of accumulated damage.

In line with stochastic, physics-inspired models (e.g., Langevin-type descriptions with a noise power *D*_0_), aging in short-lived, unstable animals can be formalized as a first-passage problem on an unstable landscape, where interventions that reduce instability or raise the barrier to failure “rescue” animals close to a death threshold without resetting prior damage. This framing explicitly connects our work to non-equilibrium statistical mechanics and complex systems theory. The model described here constitutes a limiting case within a broader class of dynamical systems that also includes “stable” organisms. In early life, these systems (including humans) reside in a dynamically stable regime for most of their lives. Progressive entropy-driven parameter drift reduces stability over time, eventually bringing the system to a codimension-1 bifurcation, beyond which dynamics become intrinsically unstable. By contrast, short-lived organisms such as nematodes appear to operate in or rapidly transition into this unstable regime early in adulthood, possibly shortly after reproductive maturity [11].

This distinction carries a concrete clinical prediction: interventions that shift *z* dynamics (such as senolytics, anti-inflammatory agents, or proteostasis enhancers) will produce persistent effects in unstable animals like *C. elegans*, but only transient benefits in stable organisms like humans, where resilience mechanisms restore *z* toward equilibrium after the perturbation is removed. To extend the maximum lifespan of stable organisms requires slowing the rate of entropic damage accumulation or boosting the initial stability margin—targets that are mechanistically distinct from those optimal for short-lived model organisms.

### Workshop Question 1: What under-appreciated or emerging direction are you highlighting, particularly one connected to your own work?

Recent physics-inspired frameworks for aging fall into two broad classes that yield mathematically similar stochastic descriptions but interpret the underlying variables differently. Dynamical-systems models proposed by Fedichev and colleagues [10, 11, 22] treat aging as the progressive loss of stability in organism-level regulatory dynamics, with state variables interpreted as emergent collective modes representing damage and stress-responses. Saturation-removal models developed by Alon and colleagues [23, 24], attribute Gompertzian mortality to the nonlinear production-removal kinetics of specific damage-associated biological variables (for example, senescent cells or aggregated proteins).

Experiments such as the DAF-2 degron system provide a particularly stringent testbed for comparing such frameworks. Both classes of models can reproduce key features of aging trajectories, including early Gompertzian mortality, late-life deceleration of mortality increases, and apparent age-dependent intervention effects at the population level. The relevant question is therefore not whether a given phenomenon can be reproduced in principle, but which structural assumptions are required to do so, and how these map onto experimentally observable quantities.

A central observable in the degron system is the relationship between intervention timing, residual lifespan, and state-dependent outcomes. Intervention at late ages, after substantial mortality has already occurred, can restore remaining lifespan close to the maximum observed under early-life intervention [2, 3]. Interpreting this result requires disentangling chronological age, state selection, and intervention-induced shifts in the underlying dynamical variables. Different model classes attribute these effects to distinct mechanisms such as explicit time dependence in production terms, state-dependent dynamics in unstable modes, or modifications of effective thresholds, leading to different predictions for conditional survival, hazard trajectories, and late-life survival tails.

A second discriminator is the selective persistence of age-related pathology. The degron extends lifespan while leaving pharyngeal degeneration, gonadal atrophy, yolk pooling, uterine tumors, and muscle PolyQ aggregates intact, yet simultaneously restoring proteostasis, stress resistance, and cuticle integrity [3].

In the context of our framework, this heterogeneity maps onto the two-variable structure of the instability framework [11]: irreversible entropic damage *Z* accumulates in structural pathologies that the intervention does not reset, while the dynamic resilience mode *z*_0_ is modulated by the intervention and restores the features coupled to it. Whether and how this observation can be reproduced within the saturation-removal framework lies beyond the scope of the present work, but is an interesting question. More broadly, such experiments highlight the need for systematic, state-resolved comparisons between competing theoretical frameworks. More precise measurements that combine longitudinal state, high-resolution survival and mortality analysis, and perturbations at different points along the trajectory will provide a direct route to distinguishing between mechanisms that are otherwise difficult to separate at the level of aggregate survival curves alone.

### Workshop Question 2: How does this direction relate to physics—through mathematical modeling, complex systems approaches, materials science, or other relevant modalities?

We connect aging to physics by modeling *C. elegans* decline as a stochastic, Langevin-type process evolving along an intrinsically unstable dynamical mode. The noise power *D*_0_ captures the amplitude of physiological fluctuations, while deterministic drift amplifies them over time until a failure threshold is reached, resulting in death as a first-passage event. Within this stochastic dynamical framework, interventions such as late-life DAF-2 degradation are naturally interpreted as changes to the system’s instability parameter *α*: rather than restoring stability or reversing accumulated damage, the intervention reduces the growth rate of the unstable mode and shifts critical thresholds, delaying threshold crossing.

### Workshop Question 3: How many research groups are currently active in this area?

The Jan Gruber’s lab in Singapore is actively working on late-life auxin-induced degradation of DAF-2 in relation to the physical model proposed by Peter Fedichev. Previously, the labs of David Gems, Matt Kaeberlein, Clara Essmann, and Della David have collaborated with Collin Ewald’s lab on the late-life auxin-induced degradation of DAF-2 [2, 3]. Finally, the David Gems lab continues to experimentally investigate individual frailty with late-life DAF-2 degron interventions. Given that this is such an easy-to-use tool and an underexplored area of research, the aspiration is to inspire more C. elegans labs and physicists to work on late-life lifespan extension. The DAF-2 degron C. elegans are available at the Caenorhabditis Genetics Center [25].

### Workshop Question 4: What key elements are still lacking—such as specialized workshops, improved instrumentation, additional measurements, or new modeling approaches?

We lack standardized longitudinal datasets explicitly designed to test dynamical models, including repeated measures of resilience, proteostasis capacity, and other putative control variables across the lifespan under defined perturbations. Also, no model is perfect. However, it is striking that this Langevin-type model captures late-life longevity dynamics without explicitly parameterizing damage or waste accumulation. Remarkably, however, this simple Langevin-type framework can explain why interventions such as late-life DAF-2 degradation extend lifespan even when applied to very old animals. In this model, the organism does not “know” its chronological age. The probability of death depends only on its position along the dynamical failure coordinate that governs the approach to the death threshold.

Animals occupying the same position along this coordinate are therefore equivalent with respect to mortality dynamics, regardless of when that state is reached. Genetic or environmental interventions act by changing the dynamical parameters that govern motion along this co-ordinate. Because the intervention modifies the dynamics themselves rather than reversing accumulated damage, it has the same effect in young and old animals that occupy comparable positions in this state space. This contrasts with damage-driven frameworks in which aging reflects the progressive accumulation of defects over time, making the response to intervention inherently age-dependent.

Although many senescent pathologies—such as pharyngeal degradation, gonadal atrophy, intestinal atrophy, increased yolk and uterine tumors—were not repaired nor rejuvenated upon late-life DAF-2 degradation, these animals still experienced a doubling of their lifespan [3].

Whereas the decline in the structural integrity of their exoskeleton (cuticle) was markedly slowed, possibly by improving protein homeostasis [3]. Other pathologies, such as age-dependent protein aggregates inside and outside cells, were removed upon late-life DAF-2 degradation, suggesting rejuvenation through improved protein homeostasis [3]. However, not all protein aggregates were removed. For instance, the Huntington disease PolyQ that age-dependently aggregates in *C. elegans* muscles persisted even when the animal’s lifespan was doubled by late-life DAF-2 degradation [3]. This might suggest that it is not an organism-wide rejuvenation of proteostasis but rather is targeted to “life-threatening” damage or waste products. Thus, this suggests that longevity is governed by the state-space dynamics of a few critical ‘life-threatening’ variables rather than a global reversal of all age-related damage. Future research must therefore focus on identifying the specific control variables that dictate the system’s proximity to the death threshold.

### Workshop Question 5: What would be required to accelerate progress in this field (e.g., the level and structure of funding, duration, and whether support should come from academia or industry)?

Progress would be accelerated by physicists joining the aging field to build models, similar to this one here, that could explain the underlying molecular mechanisms of aging and how to connect these fast-acting processes, such as chemical reactions, with the decades of decline of physiological functions observed during aging. From patchy, imperfect longitudinal molecular data, can we determine tipping points into disease? Or would cross-sectional cohort data suffice, given that, during human aging, sharp tipping points have already been observed in several omics datasets [26–28]? Because this is basic, fundamental research, the funding should come from academia and foster more interdisciplinary collaboration between physicists and biologists. From governments, industry, and consortia, there needs to be more investment in large cohort data and open-access sharing to study human aging biology.

### Workshop Question 6: What contributions could industry make—for example, developing better instruments or enabling large-scale data collection?

Industry should consider sharing large internal human clinical datasets in a joint, non-competitive space, as it has previously done for Alzheimer’s disease [29, 30]. For instance, only the placebo-controlled arm of clinical trials could be shared (without infringing on intellectual property) to see whether it is possible to build a large synthetic aging cohort. Industry could provide expertise and infrastructure for large-scale data engineering and analysis, enabling the collection and integration of the dense longitudinal datasets needed to fit and validate dynamical models of aging.

### Workshop Question 7: Are there regulatory barriers that should be reconsidered, and in which countries?

There are important regulatory and governance gaps around access to human clinical and cohort data. In some countries, restrictive rules limit the use and sharing of clinical trial data, while in others, there is still no clear framework mandating or enabling responsible open access to large cohort resources (such as the UK Biobank). Moreover, entrenched academic incentives often allow senior investigators to control access to cohorts for many years, prioritizing their own publication pipelines over broader community use.

### Workshop Question 8: What potential “unknown unknowns” might exist: new interventions, model organisms, or methodological approaches that could fundamentally shift the field?

Beyond these questions, there are likely deeper “unknown unknowns” that only become apparent once we push our current models to their limits. For example, even if a simple Langevin-type instability can describe aging in short-lived, unstable organisms, we still do not know whether this is merely a convenient coarse-graining of a much higher-dimensional biochemical network or whether there really is a small set of effective variables that governs organismal decline across systems. Can we actually connect fast microscopic events, such as chemical reactions, protein misfolding, or turnover of damaged components, to the slow deterioration over years or decades, rather than just introducing ad-hoc fitting parameters into our models to cover up what we do not yet understand in detail? In that light, what is aging: the emergence of an unstable dynamical mode, a slow entropy-driven drift of control parameters, or something more akin to an effective “force/gravity” in physiological state space that steadily pulls organisms away from health toward disintegration? And, if so, is it possible to redirect this force rather than only counteract its consequences? Cases such as the so-called “immortal” hydra, which exhibit negligible senescence at the organismal level, suggest that some life histories may operate in a regime in which the failure threshold is continually repaired or never reached, implying that a fixed death boundary may not be universal. This in turn raises a more radical possibility: are there interventions that do not primarily act on damage or single pathways, but instead change the shape or boundaries of the underlying regulatory landscape itself, and if such interventions exist, what kinds of model organisms, measurements, and theoretical tools would be needed to even recognize them when we encounter them?

## MATERIALS AND METHODS

The *C. elegans* strains and experiments are described in [2, 3]. The DAF-2degron *C. elegans* is available at https://cgc.umn.edu/strain/RAF2181.

Organismal deterioration was modeled as a one-dimensional stochastic process following the instability framework described by Podolskiy et al. [10]. We defined death as a first-passage event, occurring when the dynamical trajectory *z*(*t*) first reached a critical failure threshold *Z* = *α/g*. All trajectories were initialized at *z* = 0, representing the baseline healthy state.

The system dynamics were governed by the Langevin equation

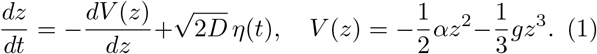

We integrated this equation using the Euler–Maruyama method with a fixed time step Δ*t* = 0.05.

To simulate late-life *daf-2* auxin intervention, we implemented a piecewise parameter shift at time *t*_int_ = 21. During the pre-intervention period (*t* < *t*_int_), the system evolved with a high linear instability parameter *α*, and this parameter was reduced in the post-intervention period (*t*≥ *t*_int_). Crucially, the state variable *z* remained continuous during the whole duration, and the diffusion coefficient *D* was fixed. This ensured the intervention modified the future stability landscape without resetting accumulated physiological deterioration.

Simulations were implemented in Python 3.12. Numerical computations were performed using NumPy, and survival data were visualized using Matplotlib and Seaborn.

## AUTHOR CONTRIBUTIONS

All authors participated in analyzing and interpreting the data. C.Y.E. and J.G. designed the experiments. D.L. performed *in silico* analysis and modeling of lifespan assays. All authors contributed to writing the manuscript.

## CONFLICTS OF INTEREST

The authors declare that the research was conducted in the absence of any commercial or financial relationships that could be construed as a potential conflict of interest. With no relation to the present manuscript, C.Y.E. declares to be a co-founder and shareholder of Avea Life AG and Lichi3 GmbH and is employed by Novartis. A.M. is employed by Novartis. P.F. is a shareholder and employee of Gero AI. J.G. declares that he is co-founder and shareholder of Antikythera Laboratories Pte. Ltd.

## ACKNOWLEDGMENTS

We thank WormBase for curated gene and phenotype information and the Caenorhabditis Genetics Center (CGC), which is funded by NIH Office of Research Infrastructure Programs (P40 OD010440), for storing and distributing the DAF-2degron strain.

## Notes

### Competing Interest Statement

The authors have declared no competing interest.

